# Damage to the Right Insula Disrupts the Perception of Affective Touch

**DOI:** 10.1101/592014

**Authors:** Louise P. Kirsch, Sahba Besharati, Christina Papadaki, Laura Crucianelli, Sara Bertagnoli, Nick Ward, Valentina Moro, Paul M. Jenkinson, Aikaterini Fotopoulou

## Abstract

Specific, peripheral C-tactile afferents contribute to the perception of tactile pleasure, but the brain areas involved in their processing remain debated. We report the first human lesion study on the perception of C-tactile touch (N = 59), revealing that posterior and anterior right insula lesions reduce tactile, contralateral and ipsilateral pleasantness sensitivity, respectively. These findings are consistent with a posterior-to-anterior pattern of integration of interoceptive information in the frontoinsular junction.

## Introduction

Increasing evidence points to the importance of affective touch to human development and health (McGlone et al., 2014). Humans may have a specialized neurophysiological system for tactile affectivity, separate from that for touch discrimination. Specifically, in the peripheral nervous system, affectivity of touch has been linked to the activation of unmyelinated, mechanosensitive C-tactile fibers (CTs) that are present only in human hairy skin and respond preferentially to low pressure, slow stroking touch at skin temperature (Löken et al., 2009; Ackerley et al., 2014). Microneurography studies found that CTs are velocity tuned, responding optimally to a stimulus moving over their receptive field at between 1 and 10cm/s, with discharge frequencies that correlate with subjective ratings of stimulus pleasantness as measured psychophysically (Löken et al., 2009). Functional neuroimaging studies have demonstrated a functional segregation, with primary and secondary somatosensory cortices most commonly associated with discriminatory touch (Aβ mediated), while tactile pleasantness (CT mediated) is associated with other areas such as the posterior insula (Morrison, 2016), parietal operculum, orbitofrontal cortex and superior temporal sulcus (Perini et al., 2015; Björndotter, 2016). These studies however are correlational. Only two neuromodulatory, repetitive transcranial magnetic stimulation (rTMS) studies (Case et al., 2016; 2017) have investigated causal relationships, finding that the right primary and secondary somatosensory cortex are not necessary for the perceived affectivity of touch. The causative role of the insular cortex, subcortical structures and white matter connections has not yet been studied, as virtual lesion methods have limited validity when applied to these deeper regions (Lenoir et al., 2018). By contrast, lesion studies allow for direct examination of the functional role of both superficial and deep brain areas. Accordingly, we aimed to investigate for the first time the right hemisphere regions which are necessary for the perceived affectivity of CT-optimal touch, applying a voxel-based lesion symptom mapping approach^10^ (VLSM; Bates et al., 2003) in a large, consecutively recruited cohort of patients (N=59) with recent, first-ever, right hemisphere lesions following a stroke. The selection of right-hemisphere patients restricts any laterality interpretations, but it also avoids the possibly confounding sequelae of left hemisphere lesions, such as language and depression symptoms (Robinson et al., 1984; Whyte et al., 2002).

We used a previously validated tactile stimulation paradigm (Crucianelli et al., 2013; 2018; Gentsch et al., 2015; von Mohr et al., 2017; Kirsch et al., 2018), together with standardized neuropsychological, somatosensory and mood assessments. Our affective touch paradigm required blindfolded patients (N=59, RH) and age-matched healthy controls (N=20, HC), to rate the intensity and pleasantness of brushing stimuli delivered at velocities known to activate the CT-system optimally (3cm/s; CT-optimal affective touch) or not (18cm/s; CT-suboptimal neutral touch) to both the left (contralesional) and the right (ipsilesional) forearm (see Methods). Participants had to rate also the hypothetical pleasantness of being touched by a typically pleasant vs. unpleasant material.

Given that right hemisphere and particularly right perisylvian regions have been previously associated with somatosensory and interoceptive perception (Dijkerman & de Haan, 2007; Preusser et al., 2015), we expected our patients to have, on average, reduced ratings of both touch intensity and pleasantness in comparison to healthy controls, and particularly in the contralesional left forearm. Crucially, given the assumed neurophysiological specificity of the CT system, we expected that more specific lesions to the posterior insula (Morrison, 2016) would lead to a lack of CT pleasantness sensitivity (defined as the pleasantness difference between CT-optimal and CT-suboptimal velocities), particularly on the contralesional forearm.

## Methods

### 1. Subjects and clinical investigation

Fifty-nine, unilateral, right-hemisphere-lesioned stroke patients (mean age: 65.86 ± 14.12 years; age range: 38-88 years; 31 females) were recruited from consecutive admissions to seven stroke wards as part of a larger study using the following inclusion criteria: (i) imaging-confirmed first ever right hemisphere lesion; (ii) contralateral hemiplegia; (iii) <4 months from symptom onset; (iv) no previous history of neurological or psychiatric illness; (v) >7 years of education; (vi) no medication with significant cognitive or mood side-effects; (vii) no language impairments that precluded completion of the study assessments; and (viii) right handed. All participants gave written, informed consent to take part in the study. The local National Health System Ethics Committees approved the study, which was carried out in accordance to the Declaration of Helsinki.

All patients underwent a thorough neurological and neuropsychological examination. Premorbid intelligence was assessed using the Wechsler Test of Adult reading (WTAR; Wechsler, 2001). Post-morbid, general cognitive functioning, including long-term verbal recall was assessed using the Montreal Cognitive Assessment (MoCA; Nasreddine, 2005). The Medical Research Council scale (MRC; Guarantors of Brain, 1986) was used to assess limb motor strength. Proprioception was assessed with eyes closed by applying small, vertical, controlled movements to three joints (middle finger, wrist and elbow), at four time intervals (correct = 1; incorrect = 0; Vocat et al., 2010). Working memory was assessed using the digit span task from the Wechsler Adult Intelligence Scale III (Wechsler, 1997). The Hospital Depression and Anxiety Scale (HADS; Zigmind and Snaith, 1983) was used to assess anxiety and depression. Executive and reasoning abilities were assessed using the Frontal Assessment Battery (FAB; Dubois et al., 2000). Four subtests of the Behavioural Inattention Test (BIT; Wilson et al., 1987) were used to assess visuospatial neglect. Personal neglect was assessed using the ‘one item test’ (Bisiach et al., 1986) and the ‘comb/razor’ test (Mcintosh et al., 2000).

Twenty age-matched healthy control subjects were recruited and tested with the same behavioural paradigm in order to assess the specificity of deficits in the patient group (healthy control group; 63.05 ± 12.12 years; age range: 46-87 years; 11 females). Patients’ demographic characteristics and their performance on standardized neuropsychological tests and how they compared to the healthy sample are summarized in Table 1.

**Table 1.** Summary of demographics and neuropsychological data. Description: Nottingham**^a^** = Light Touch subscale of the Revised Nottingham Sensory Assessment^31^ (rNSA; Lincoln, Jackson, & Adams, 1998; score overall for each arm with 0: no sensation; 1: slightly impaired; 2: no deficit); MRC = Medical Research Council scale (Guarantors of Brain, 1986); MOCA = The Montreal Cognitive Assessment (Nasreddine et al., 2005); FAB = Frontal Assessment Battery (Dubois et al., 2000); Premorbid IQ-WTAR= Wechsler Test of Adult Reading (Wechsler, 2001); HADS = Hospital Anxiety and Depression scale (Zigmond & Snaith, 1983); Comb/razor test = tests of personal neglect (MacIntoch, Brodie, & Beschin, 2000); Bisiach one item test = test of personal neglect; line crossing, star cancellation, copy & representational drawing = conventional sub-tests of Behavioural Inattention Test (Wilson, Cockborn & Halligan, 1987). Dashed line indicates not applicable. Due to several clinical constraints (e.g. fatigue, acceptance and time constraints), we have a number of missing data on these tests. Specific numbers are indicated in the right column. N_RH_=number of right hemisphere stroke patients having fully completed the corresponding test. N_HC_= number of healthy controls having fully completed the corresponding test. ^*^ Significant difference between groups, p < .05.

|  | Stroke Patients –<br>RH (N=59; 31<br>females) |  | Healthy Controls<br>- HC (N=20, 11<br>females) |  | Mann-Whitney Test | N<sub>RH</sub>/N<sub>HC</sub> |
| --- | --- | --- | --- | --- | --- | --- |
|  | Mean | SD | Mean | SD |  |  |
| Age (years) | 65.86 | 13.87 | 63.05 | 12.12 | $U(78)=514.00$, $Z=-.857$, $P=0.391$ | $N=59/20$ |
| Education (years) | 11.40 | 2.87 | 14.75 | 2.82 | $U(70)=211.50$, $Z=-3.906$, $P<0.001^*$ | $N=52/20$ |
| Days from onset | 16.95 | 18.68 | - | - |  |  |
| Orientation | 2.80 | 0.41 | - | - |  |  |
| Nottingham<sup>a</sup> on left arm (max 2) | 0.66 | 0.78 | - | - |  |  |
| Nottingham<sup>a</sup> on right arm (max 2) | 2 | 0 | - | - |  |  |
| Proprioception (max 9) | 5.10 | 2.64 | - | - |  |  |
| MRC Left upper limb | 0.30 | 0.75 | - | - |  |  |
| Digit span forwards | 5.95 | 1.40 | 6.58 | 1.83 | $U(66)=279.50$, $Z=.936$, $P=0.349$ | $N=56/12$ |
| Digit span backwards | 3.50 | 1.55 | 4.75 | 1.28 | $U(66)=177.00$, $Z=-2.621$, $P=0.009^*$ | $N=57/12$ |
| MOCA | 19.85 | 5.18 | 28.19 | 1.92 | $U(45)=5.50$, $Z=-4.271$, $P<0.001^*$ | $N=39/8$ |
| MOCA memory subscale | 2.92 | 1.78 | 4.00 | 1.60 | $U(45)=95.00$, $Z=-1.769$, $P=0.077$ | $N=39/8$ |
| Premorbid IQ-WTAR | 34.00 | 9.35 | 49.11 | 1.69 | $U(25)=3.00$, $Z=-4.037$, $P<0.001^*$ | $N=18/9$ |
| HADS depression | 5.75 | 3.49 | 3.13 | 2.19 | $U(50)=150.00$, $Z=-2.593$, $P=.010^*$ | $N=37/18$ |
| HADS anxiety | 8.02 | 4.33 | 6.06 | 3.01 | $U(50)=208.00$, $Z=-1.409$, $P=0.159$ | $N=37/18$ |
| FAB total score | 11.38 | 4.02 | - | - |  |  |
| Comb/razor test bias (%bias) | -23.37 | 27.06 | - | - |  |  |
| Bisiach one item test | 0.47 | 0.68 | - | - |  |  |
| Line crossing (max 36) | 22.56 | 11.85 | - | - |  |  |
| Star cancelation (max 54) | 29.93 | 18.23 | - | - |  |  |
| Copy | 0.87 | 1.20 | - | - |  |  |
| Representational drawing | 0.62 | 0.93 | - | - |  |  |
| Line bisection | 2.87 | 3.05 | - | - |  |  |

### 2. Design and Predictions

The present study aimed to investigate the neuroanatomical bases of affective touch. To this aim, we compared a large cohort of right hemisphere stroke patients to healthy controls, and explored how deficits in affective touch perception are linked with specific brain lesions. To achieve this aim we applied an affective touch paradigm that manipulated three factors: i) the velocity of the touch applied (slow, CT-optimal, affective touch vs. fast, CT-suboptimal, neutral touch); ii) the forearm the touch was applied to (right, ipsilesional vs. left, contralesional); iii) and the group of participant (Stroke patients vs. Healthy controls). For each type of touch we recorded two measures: 1) intensity ratings and 2) pleasantness ratings. We also asked participants to rate the pleasantness of imagined touch with either a smooth material versus a rough material, to control for top-down effects.

To investigate the neuroanatomical bases of affective touch, we conducted two main voxel-based, lesion-symptom mapping analyses, separately for each forearm, using as predictors the CT pleasantness sensitivity (difference between average pleasantness ratings for CT-optimal touch and CT-suboptimal touch). In addition to the main analyses we also ran a control analysis, using the difference between hypothetical pleasantness ratings of pleasant (velvet) and unpleasant (sandpaper) material as predictors, to control for potential top-down affective deficit. Finally, a lesion overlap was calculated to create a color-coded overlay map of lesioned voxels across participants with negative or null CT pleasantness sensitivity on each forearm.

Given our patients’ lesions to several perisylvian regions of the right hemisphere previously associated with somatosensory perception (Dijkerman & de Haan, 2007; Preusser et al., 2015), we expected that our patients would have, on average, reduced ratings of both touch intensity and pleasantness in comparison to healthy controls, and specifically in the contralesional left forearm. However, we did not expect a general right stroke effect on pleasantness sensitivity to CT affective touch (defined as the pleasantness difference between CT-optimal and CT-suboptimal velocities), given the assumed neurophysiological specificity of the CT system. Instead, we expected that lesions involving mainly the right posterior insula (Morrison, 2016) would lead to a lack of pleasantness sensitivity, particularly on the contralesional forearm. Moreover, as some authors have proposed that the right hemisphere, and particularly the right anterior insula, has a crucial role in interoceptive awareness for the entire body (Craig, 2009; Critchley et al., 2004; Kann et al., 2016; Khalsa et al., 2009; Salomon et al., 2016), we expected also to find some causal role of ipsilateral areas (right hemisphere regions after touch on the right forearm) and particularly the right anterior insula in the perception of affective touch on the ipsilesional forearm.

While testing only right-hemisphere patients restricts the conclusions of the study to the right hemisphere (as in the aforementioned neuromodulatory studies; Case et al., 2016, 2017) and hence our conclusions regarding laterality are preliminary, it avoids confounds related to the many cognitive and emotional sequel of left hemisphere lesions, such as language problems and depressive reactions (Robinson et al., 1984; Whyte et al., 2002).

### 3. Affective touch protocol

Tactile stimulation followed a previously validated protocol (Crucianelli et al., 2013; 2018; Gentsch et al., 2015; von Mohr et al., 2017; Kirsch et al., 2018), including both ‘imagined’ and actual touch questions. Specifically, first a 9cm × 4cm area of skin stimulation was marked on both forearms and patients were familiarized with the vertical rating scales (to minimize the effects of neglect; we also always ensured the participants could see the scale and read it aloud to facilitate them), anchored at “0 - not at all” and “10 - extremely”. We first sampled top-down, prior beliefs about tactile pleasantness by asking two hypothetical questions about imagined touch: “How pleasant would it be to be touched by velvet on your skin” (typically pleasant) and “How pleasant would it be to be touched by sandpaper on your skin?” (typically unpleasant). Participants were asked to answer using the vertical 0 to 10 pleasantness scale.

We then explained that actual tactile stimuli would be delivered on the marked forearm areas, while participants were blindfolded, and instructed to remain still and to focus on both the intensity and pleasantness of the touch they were experiencing (Fig. 1). Tactile stimuli were administrated by a 4 cm wide soft make up brush made from natural hair (Natural hair Blush Brush, No. 7, The Boots Company). Brush strokes were administered by a trained female experimenter in proximal-to-distal direction with the brush held in a perpendicular position, with the edges of the brush tracking the width of the testing area to control for pressure. Every touch condition lasted for 3 seconds; with an inter-stimuli interval of at least 30s. After each touch, participants were asked to answer two questions: first “How well did you feel the touch?” (i.e. touch intensity rating), and if they felt the touch (i.e. reporting an intensity rating >0), they were asked “How pleasant was the touch?” (i.e. touch pleasantness rating), using the above described 0 to 10 vertical scale. Tactile stimuli were delivered at two different velocities on the participant’s left and right forearm: CT-optimal speed (3cm/s, known to activate CT fibers optimally; one stroke over the 9cm long area) and CT-suboptimal speed (18cm/s, known to activate CT fibers to a lesser degree, suboptimally^;^ Gentsch et al., 2015; six strokes). Each condition was repeated 6 times, leading to a total of 24 trials – delivered in a pseudorandomized order. The experiment was split into 3 blocks to avoid fatigue; short breaks were taken after a set of 8 trials (2 repetitions of each condition in each block).

**Figure 1.**
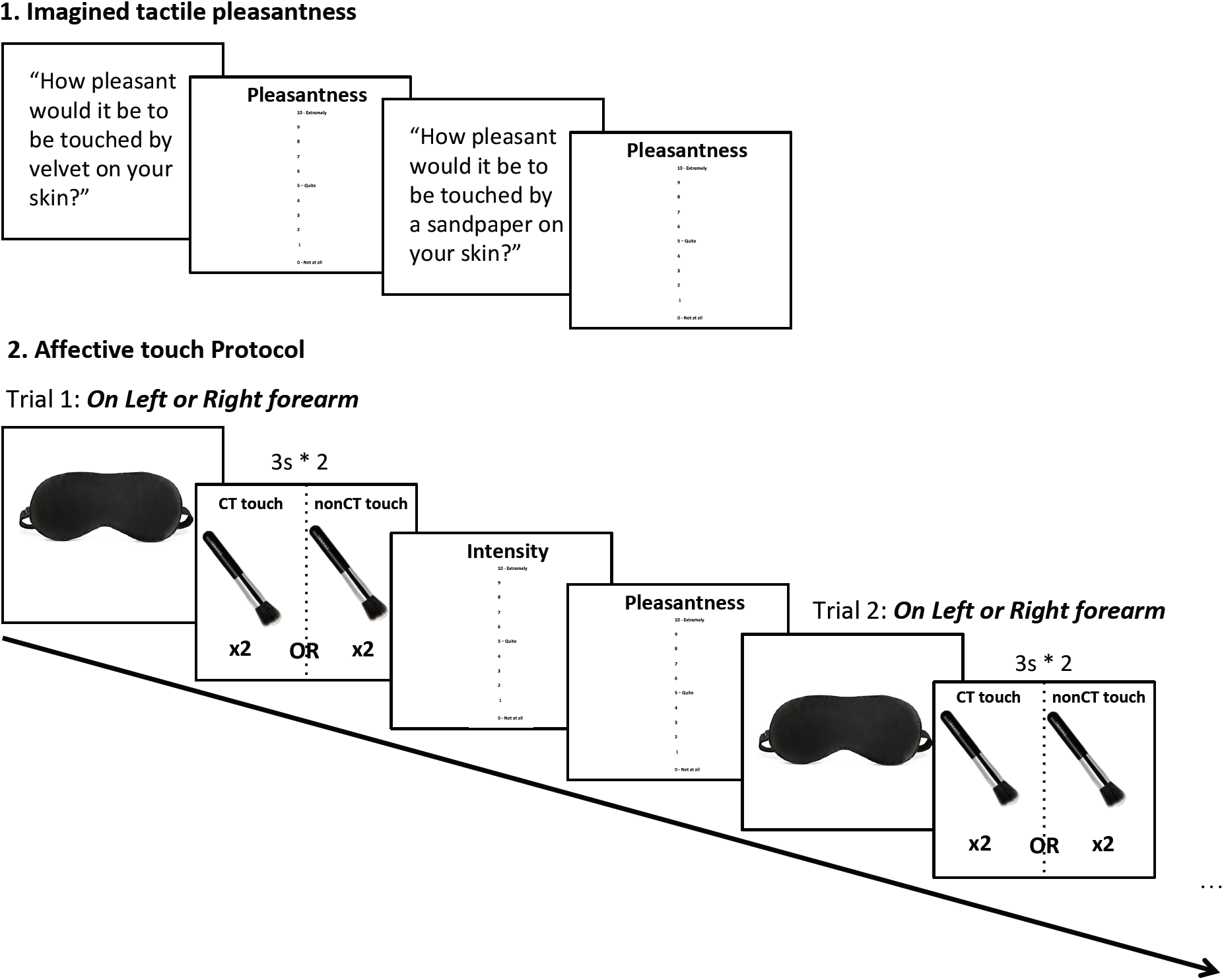
Experimental design and timeline. 1. Participants were first asked to answer two hypothetical questions about imagined touch: “How pleasant would it be to be touched by velvet on your skin” (typically pleasant) and “How pleasant would it be to be touched by sandpaper on your skin?” (typically unpleasant). Participants were asked to answer using the vertical 0 to 10 pleasantness scale. 2. Participants were then asked to put on a blindfold at the onset of each trial before the experimenter delivered the touch on the left or right forearm at CT-optimal or CT-suboptimal velocities (pseudorandomized), each touch lasted for 3 seconds and was repeated twice with a one second break in between. After each touch, blindfold was removed so participants could rate the touch on two scales: Intensity = How well they felt the touch; and Pleasantness = How pleasant was the touch, each on a vertical scale ranging from 0, not at all, to 10, extremely. After ratings were recorded, the participant was asked to put the blindfold back before starting the next trial.

All patients had intact sensation on the right ipsilesional forearm (i.e. rated the intensity of tactile stimuli as greater than zero in all the trials, irrespective of velocity, and had intact sensation on this side according to a standardized assessment; the Revised Nottingham Sensory Assessment (rNSA; Lincoln, Jackson, & Adams, 1998) but as predicted, on the contralesional side, some patients (40.7%, N=24) were not able to perceive the tactile stimuli (corroborated also by the above standardized somatosensory assessment), and therefore gave a rating of zero on the intensity scale, and were not asked to provide pleasantness ratings. Thus, pleasantness ratings were available only from the remaining 35 patients who were able to perceive the intensity and pleasantness of most contralesional tactile stimuli in our paradigm.

### 4. Lesion mapping methods and analyses

Routinely acquired clinical scans obtained on admission (<2 days post stroke) were collected for the 59 patients (49 via computerized tomography, CT; and 10 via magnetic resonance imaging, MRI). We note that testing patients in the acute post-stroke phase entails challenges but avoids any confounds relating to plasticity and functional reorganization (Baier et al., 2014; De Haan & Karnath, 2018). The patient’s lesion was mapped by means of the MRIcron software (Rorden & Brett, 2000) on the standard T1-weighted MRI template (ICBM152) of the Montreal Neurological Institute (MNI) coordinate system. Lesions from these scans were segmented and co-registered using a manual procedure, as this method remains the best methods to date for lesion mapping of clinical scans and shown to be more accurate than automatized methods (Maier et al., 2015; de Haan & Karnath, 2017; Liew et al., 2018). Two expert clinicians, blind to the hypotheses of the study, outlined the lesions. In the case of disagreement of two lesion plots, the opinion of a third, expert anatomist was requested. Scans were registered to the T1-weighted MRI scan template (ICBM152) of the Montreal Neurological Institute, furnished with the MRIcron software (ch2, http://www.cabiatl.com/mricro/mricron/index.html). First, the standard template (size: 181 × 217 × 181 mm, voxel resolution: 1mm^2^) was rotated on the three planes in order to match the orientation of the patient’s MRI or CT scan. Lesions were outlined on the axial slices of the rotated template. The resulting lesion volumes were then rotated back into the canonical orientation, in order to align the lesion volumes of each patient to the same stereotaxic space. Finally, in order to exclude voxels of lesions outside white and gray matter brain tissue, lesion volumes were filtered by means of custom masks based on the ICBM152 template.

The statistical contribution of lesion location to CT pleasantness sensitivity and fabrics deficits was tested using voxel-based lesion symptom mapping (VLSM), and using the behavioral scores as continuous predictor. The statistical process performed in voxel-based lesion–symptom mapping (Bates et al., 2003) consists of the following steps: at each voxel of the spatially standardized scan images, patients are divided into two groups according to whether they did or did not have a lesion affecting that voxel. Behavioral scores are then compared for these two groups with a t-test, yielding a single-tailed p-value for each voxel^1^. This method allows controlling for lesion size, which is included as a covariate of non-interest. Note that to avoid spurious results due to low numbers of lesioned voxels, only voxels lesioned in at least 10 participants were tested. This results in color-coded VLSM maps that represent voxels where patients with lesions show a significantly different behavioral score from those whose lesions spared that voxel at an α level of 0.01 after correction for multiple comparisons using the false discovery rate (Curran-Everett, 2000). Software to perform VLSM (http://crl.ucsd.edu/vlsm) was run using Matlab (Mathworks, 2002).

Each analysis was conducted separately for the contra- and the ipsilesional forearm, and only regions of more than 10 voxels that passed the set 0.01 FDR-corrected threshold were considered in the discussion. VLSM results were visualized in MRIcron. Three anatomical templates served to identify grey and white matter region labels: the “automated anatomical labelling” (AAL) template (Tzourio-Mazoyer et al., 2002), the JHU white-matter tractography atlas, (Mori, Wakana, van Zijl, & Nagae-Poetscher, 2005), and the “NatBrainLab” template of the “tractography based Atlas of human brain connections Projection Network” (Natbrainlab, Neuroanatomy and Tractography Laboratory; Catani & Thiebaut de Schotten, 2012; Thiebaut de Schotten et al., 2011).

## Results

### Touch perception after right hemisphere stroke - Behavioral Data

First, we investigated the effect of right hemisphere lesions on the perception of touch intensity and pleasantness, on the contralesional and ipsilesional forearm separately, by comparing stroke patients and healthy controls intensity and pleasantness ratings in turn. As the data were normally distributed, separate ANOVAs were run with touch type (CT-optimal vs. CT-suboptimal) and group (stroke patient vs. healthy controls) as factors, for each rating type and each forearm.

Given our patients’ lesions to several perisylvian regions of the right hemisphere previously associated with somatosensory perception (Dijkerman & de Haan, 2007; Preusser et al., 2015), we expected that our patients would have, on average, reduced ratings of both touch intensity and pleasantness in comparison to healthy controls, and particularly in the contralesional left forearm. However, we did not expect a general stroke effect on pleasantness sensitivity to CT-optimal touch (defined as the pleasantness difference between CT-optimal and CT-suboptimal velocities). Instead, we expected that particular lesion patterns involving the insular cortex would lead to a lack of pleasantness sensitivity, particularly on the contralesional forearm (see Morrison, 2016, for a meta-analysis).

### Touch Intensity Ratings

We were able to collect contralesional touch intensity ratings on only 39 out of the total sample of 59 patients due to an administrative error (the experimenter took binary, ‘yes’ or ‘no’ responses to the tactile stimuli instead of using the rating scale in the remaining patients). For the same reason, we only had ipsilesional touch intensity ratings for CT-optimal touch on 36 and CT-suboptimal touch on 20 patients. This unfortunately meant that our sample was reduced to 39 patients for the analyses of intensity ratings on the contralesional forearm and of 20 patients for the ipsilesional forearm.

A main effect of group on intensity ratings was observed only in the contralesional forearm, with stroke patients perceiving touch as less intense than healthy controls only on the contralesional left forearm (contralesional: F(1,57)=55.918, p<0.001, η_p_^2^ = .495; ipsilesional: F(1,38)=0.834, p=0.367, η_p_^2^ = .021); in line with the high percentage of contralesional tactile deficits in right hemisphere stroke patients (including in our patients’ sample, see Methods). Thus, even though the power of our statistical analysis was decreased on the ipsilesional side, the inability to reject the null hypothesis was consistent with our other assessments of tactile acuity for the ipsilesional side of the body (as measured by the rNSA, see Methods). We also observed a main effect of touch type on the contralesional side, with both stroke patients and healthy controls perceiving CT-optimal touch as less intense than CT-suboptimal touch (F(1,57)=4.689 p=0.035, η_p_^2^ = .076), as shown in previous studies (Löken et al., 2009; Triscoli et al., 2013). No such effect was noted in the ipsilesional side (F(1,38)=.073, p=0.789, η_p_^2^ =.002) and there was no interaction between touch type (CT-optimal vs. CT-suboptimal) and group (contralesional: F(1,57)=2.902, p=0.094, η_p_^2^ = .048; ipsilesional: F(1,38)=.954, p=0.335, η_p_^2^ = .025; see Fig. 2A&B).

**Figure 2.**
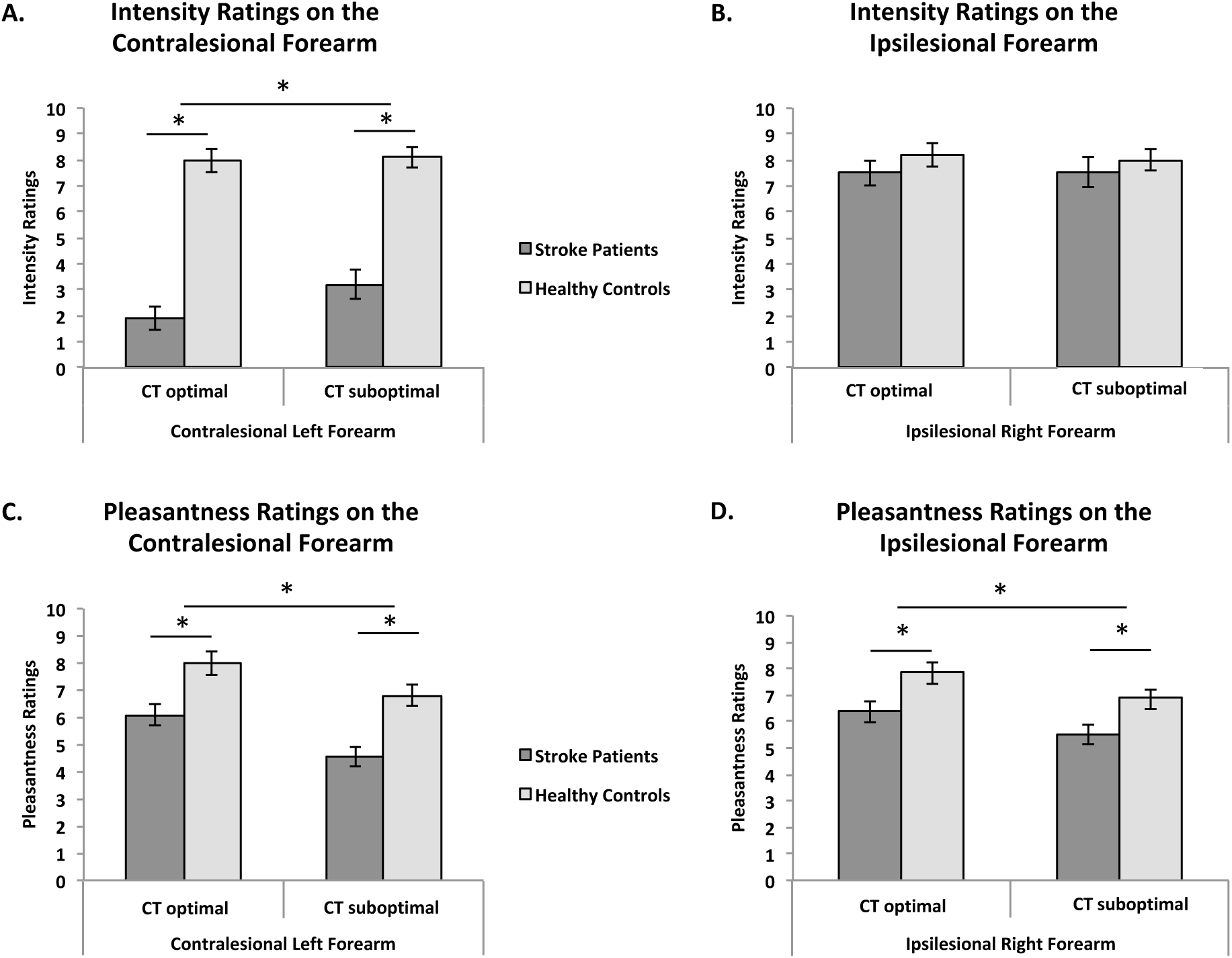
A. Average intensity ratings on the contralesional left forearm (N_RH_=39, N_HC_=20), B. Average intensity ratings on the ipsilesional right forearm (N_RH_=20, N_HC_=20), C. Average pleasantness ratings on the contralesional left forearm (N_RH_ =35, N_HC_=20), D. Average pleasantness ratings on the ipsilesional right forearm (N_RH_=41, N_HC_=20), for CT-optimal and CT suboptimal touch. Error bars represent the standard error of the mean. *depicts significant comparison, p<0.05

### Touch Pleasantness Ratings

We were able to record pleasantness ratings for contralesional forearm touch on 35 and 39 patients for CT-optimal and CT-suboptimal touch velocities respectively (data of 21 and 13 patients were missing due to the fact that patient did not feel the touch and gave an intensity rating of 0; see Methods; the remaining 3 and 8 missing data were due to administration error). For the right ipsilesional forearm, pleasantness ratings of 56 and 41 patients were recorded at CT-optimal and CT-suboptimal touch velocities respectively. Thus, the sample of the analysis of touch pleasantness was of 35 patients for the contralesional forearm and of 41 patients for the ipsilesional forearm.

Our analyses revealed a main effect of stroking type on pleasantness ratings, with CT-optimal affective touch being rated as more pleasant than CT-suboptimal neutral touch on both forearms (left contralesional: F(1,53)=22.444, p<0.001, η_p_^2^ = .297; right ipsilesional: F(1,59)=11.519, p=0.001, η_p_^2^ = .163); as well as a main effect of group, i.e. stroke patients found the touch in general less pleasant than healthy controls (contralesional: F(1,53)=14.074, p<0.001, η_p_^2^ = .210; ipsilesional: F(1,59)=7.100, p=0.010, η_p_^2^ = .107), i.e. showing a general tactile anhedonia effect. However, as expected, no interaction between touch type (CT-optimal vs. non-CT) and group was found (contralesional: F(1,53)=0.393, p=0.533, η_p_^2^ = .007; ipsilesional: F(1,59)=0.073, p=0.788, η_p_^2^ = .001; see Fig. 2C&D).

Moreover, to investigate whether the general anhedonia effect in right hemisphere patients was specific to patients that also had a tactile acuity deficit, we ran the same analysis only with patients with intact sensation (N=25), and found that the main effect of group holds, with patients finding touch overall less pleasant than controls (F(1,43)=9.880, p=0.003, η_p_^2^ = .187).

A similar general tactile anhedonia (reduced pleasantness ratings) was observed in our patients as compared to the controls for imagined tactile pleasantness in our control task (F_(1,57)_= 55.918, P<0.001, η_p_^2^=.495), but there was no Group by Fabric interaction (F_(1,70)_=.061, P=0.806, η_p_^2^=.001).

Even though as expected at a group level, stroke patients differed from healthy controls only on general tactile anhedonia, rather than CT-sensitivity (i.e. rating CT-optimal touch as more pleasant than CT-suboptimal touch), the aim of the study was to investigate the lesion patterns and neuropsychological deficits that may be associated with the inability of certain stroke patients to distinguish the pleasantness of CT-optimal versus CT-suboptimal touches. Accordingly, CT pleasantness sensitivity was calculated as the difference between the pleasantness of CT-optimal and CT-suboptimal touches. As a convention, CT pleasantness sensitivity inferior or equal to zero is considered as low in CT pleasantness sensitivity, i.e. CT affective touch perception (Crucianelli et al., 2018). On the contralesional left forearm, out of 35 patients, 10 patients (28.57%) rated CT-optimal touches as less or equally pleasant than CT-suboptimal touches; whereas on the ipsilesional right forearm, out of 41 patients, 18 patients (43.9%) rated CT-optimal touches as less or equally pleasant than CT-suboptimal touches.

Interestingly, none of the patients demographic characteristics or, neuropsychological deficits correlated significantly with their CT pleasantness sensitivity, including education, anxiety and depression scores, as well as memory as measured by the MOCA memory subscale, and premorbid intelligence, all p>0.01, alpha corrected for multiple comparisons. Thus, low CT touch sensitivity was not explained by other general cognitive deficits, as assessed in the present study. Moreover, there was no correlation between CT pleasantness sensitivities on both forearms and tactile anhedonia (as measure by the difference between the imagined pleasantness of pleasant and unpleasant fabric; r_31_=-.084 p=0.652 for the contralesional forearm, r_36_=-.086, p=.618), nor with tactile acuity as measured by intensity ratings.

### Brain regions necessary to perceiving affective touch – VLSM analyses CT pleasantness sensitivity

We investigated how deficits in ***CT pleasantness sensitivity on the contralesional forearm*** were linked to specific brain lesions. To do so, difference ratings between CT-optimal and CT-suboptimal touch on the *contralesional left* forearm were entered as continuous predictor in a VLSM analysis, with lesion volume entered as a covariate of non-interest (N=35). As shown in Table 2A, when taking into account any patients that had given pleasantness ratings on the left forearm (i.e. given intensity ratings above 1 in most trials), the only region that was identified as necessary for CT affective sensitivity on the contralesional forearm were the rolandic operculum (Fig. 3A). Subcortically, lesions extended to the tracts of the superior corona radiate. According to the white matter atlas of the Natbrainlab laboratory (Catani & Thiebaut de Schotten, 2012; Thiebaut de Schotten et al., 2011), significant voxels were found on the cortico-spinal tract, the corpus collosum, the internal capsule, and the anterior segment of the arcuate fasciculus. Moreover, including only patients without sensory deficit (participants that rated all the trials as more intense than 2; N=25) lead to lesions as well in the rolandic operculum but also, and most interestingly a bigger cluster in the posterior part of the insula (see Table 2B and Fig. 3B).

**Table 2.**
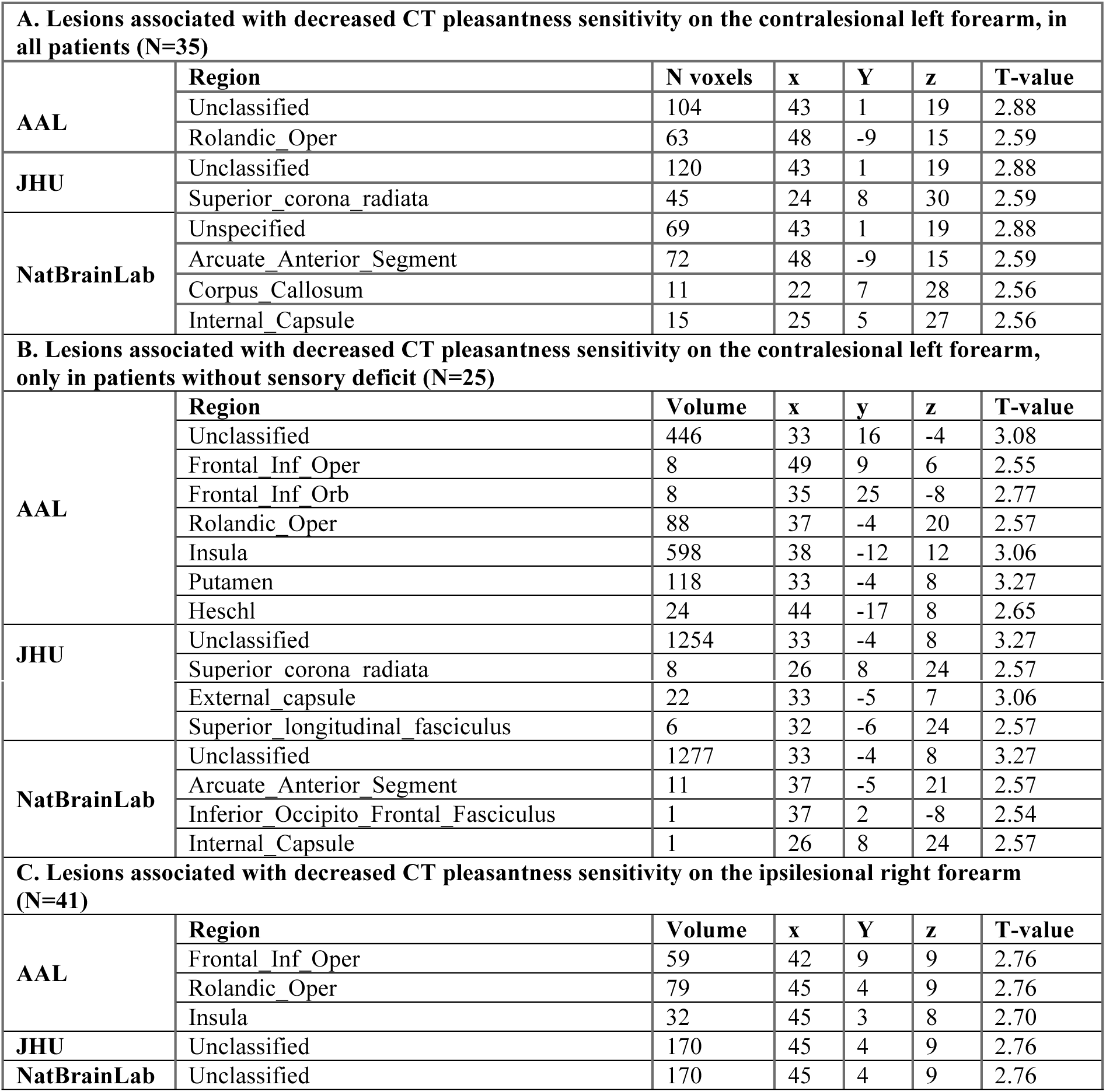
**Number of significant voxels (from the atlas of grey matter – AAL – and white matter – JHU – and NatBrainLab’s atlas) resulting from the VLSM analyses. A. with the CT pleasantness sensitivity scores for the *contralesional left* forearm as predictor**, in all patients **(N=35); B. with the CT pleasantness sensitivity scores *for the contralesional left forearm* as predictor**, only in patients without sensory deficit, N=25; **C. with the CT pleasantness sensitivity scores for the *ipsilesional right* forearm as predictor (N=41).**

**Figure 3.**
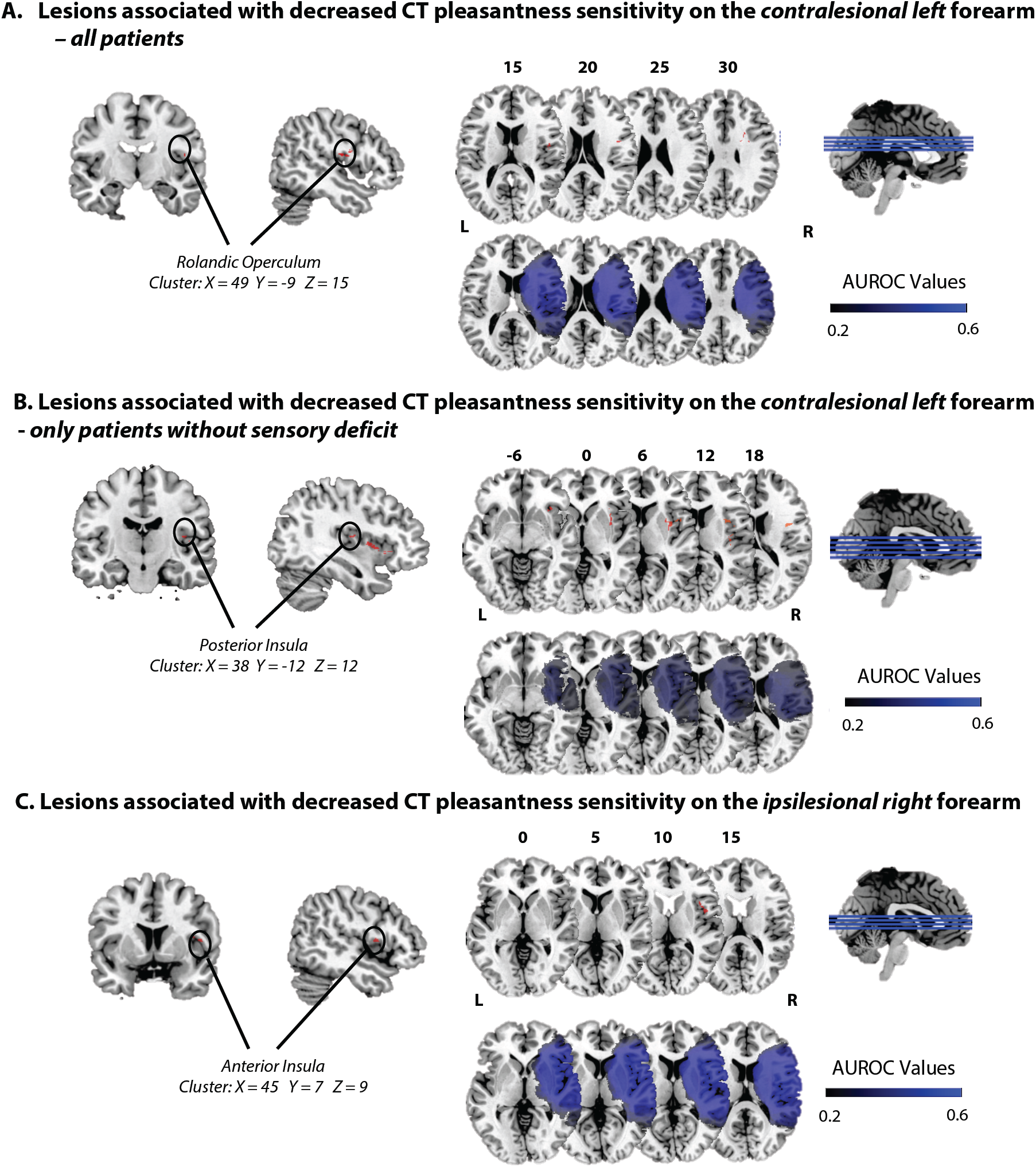
**A. Lesions associated with decreased CT pleasantness sensitivity on the contralesional left forearm,** in all patients (N=35). **B. Lesions associated with decreased CT pleasantness sensitivity on the contralesional left forearm,** only in patients without sensory deficit on the left (N=25). **C. Lesions associated with decreased CT pleasantness sensitivity on the ipsilesional right forearm** (N=41). The numbers above the brain slices indicate the corresponding MNI axial coordinates. L=Left; R=Right; The second raw represents heat maps of the voxels with power enough to detect significant results, at α=0.01, FDR-corrected. Different colors represent the area under the ROC curve (AUROC) scores, ranging from 0.2 to 0.6.

Similarly, we investigated how deficits in ***CT pleasantness sensitivity on the ipsilesional forearm*** were linked to specific brain lesions. As the perception of touch on this forearm was not affected by the stroke (as confirmed by both a standardized assessment of light touch perception – rNSA and our intensity touch ratings; see above and Methods), results of this analysis would be even more clearly associated with CT pleasantness sensitivity, albeit on the ipsilesional part of the body. To this aim, the difference between pleasantness ratings of CT-optimal and CT-suboptimal touch on the *ispilesional right* forearm (intact sensation) was entered as continuous predictor, with lesion volume entered as a covariate of non-interest (N=41). Only three regions survived the set threshold: the anterior insula, the rolandic operculum, and the frontal inferior operculum (Table 2C and Fig. 3C). This result was limited to the grey matter, as no classified regions were found with the JHU or the Natbrainlab atlases. It thus appears that the anterior insula, as well as the rolandic and frontal operculum need to be intact, even in the ipsilesional hemisphere, for individuals to perceive a pleasantness difference between CT-optimal and CT-suboptimal touch.

It is to note, that out of the 10 patients that showed a negative CT pleasantness sensitivity on the *contralesional* forearm (among all patients, N=35), 8 of them had a lesion to the rolandic operculum cluster (X=48, Y=-9, Z=15); and for the 2 remaining patients, one had a more focal deep lesions (amygdala, putamen, thalamus), that could still be on the posterior insula track; and the other had an insula lesion but more frontal; see Supplementary Figure 2A. Moreover, when taking into account only patients without sensory deficit, out of the 6 patients that showed a negative CT pleasantness sensitivity on the contralesional forearm (among patients without sensory deficit, N=25), 5 had a lesion to the posterior insula cluster (X=38, Y=-12, Z=12), and the other had a more focal deep lesion (amygdala, putamen, thalamus); see Supplementary Figure 2B.

Out of the 18 patients with a negative CT pleasantness sensitivity on the *ipsilesional* forearm, 15 had a lesion of the anterior insula cluster (X=45, Y=3, Z=8); and the three remaining had a lesion to the insula, but not on that specific cluster (see Supplementary Fig. 1C).

### Imagined Pleasantness of touch

Moreover, as control for a general pleasantness deficit, patients rated how pleasant it would be to be touched by a typically pleasant material (i.e. velvet, M_pleasantness rating_ = 6.91, SD = 1.88) and a typically unpleasant fabric (i.e. sandpaper, M_pleasantness rating_ = 0.33, SD = 0.93). Similarly as for CT pleasantness sensitivity, top-down tactile pleasantness sensitivity was computed as the difference between pleasant and unpleasant pleasantness ratings, for each patient. We considered the same patients as for the CT pleasantness sensitivity VLSM analysis (N=36 as we had missing data for 5 of them), and ran a VLSM analysis with these top-down tactile pleasantness sensitivity. This gave rise to significant voxels in the Caudate, Thalamus, Putamen and Pallidum, but crucially, not the insula (Supplementary Table 1).

## Discussion

In the present study, we used a previously validated affective touch protocol in stroke patients to investigate for the first time the right hemisphere regions, which are necessary for the perceived affectivity of CT-optimal touch, applying a voxel-based lesion symptom mapping approach. Lesion mapping results confirmed our predictions, with VLSM results corroborating the importance of the posterior insula and the rolandic operculum in perceiving CT-optimal touch on the contralateral forearm as more pleasant than CT-suboptimal touch, particularly when other tactile pathways are intact. In contrast, and most interestingly, deficits in CT pleasantness sensitivity on the ipsilesional forearm were associated with lesioned voxels in the anterior part of the insula. As patients’ perception of the discriminatory, emotionally-neutral aspects of touch on the ipsilesional forearm was not affected (verified by the lack of difference in intensity ratings between healthy controls and patients, as well as patients’ performance on a standardized somatosensory assessment), and as the left insula and somatosensory cortex of these patients were intact, these results suggest that the right anterior insula has a necessary role in the CT pleasantness sensitivity, even for the ipsilateral side of the body. Additionally, a VLSM analysis with the difference of pleasantness between hypothetical pleasant (velvet) and unpleasant (sandpaper) fabric as predictor found significant voxels subcortically in the caudate, thalamus, putamen and pallidum, but crucially, not the insula, suggesting that the above results are specific to applied tactile stimuli and not more general pleasantness comparisons.

This is the first lesion study to investigate deficits in the perceived affectivity of CT-optimal touch. Our results suggest a causal role of the posterior contralateral opercular-insular cortex for the perception of CT-optimal touch as more pleasant than CT-suboptimal touch, offering support to previous, correlational, functional neuroimaging findings on the CT system (Olausson et al., 2002; Morrison et al., 2016). In addition, our findings reveal that the *right anterior* fronto-insular junction is necessary to perceive the pleasantness of CT-optimal touch as more pleasant than CT-suboptimal touch on the ipsilateral forearm. Thus, even when the left insula and somatosensory cortex are intact and hence presumably contralateral stimuli are processed in the left cortex (as also revealed by the intact detection of ipsilesional tactile stimuli in our patients), a right anterior insula lesion is enough to cause deficits in the perception of affective touch on the right forearm. Our findings address existing debates about hemispheric laterality and interoceptive awareness (Kann et al., 2016; Khalsa et al., 2009; Salomon et al., 2016), although the VLSM method has known, intrinsic limitations and we cannot exclude the possibility of the role of the left insula in affective touch perception, nor the impact of lesions of the right hemisphere in disconnecting tracts towards the left hemisphere. However, taken together, our findings support previous findings about the functional organization and role of the human insula (Craig, 2010; Cauda et al., 2011; Kurth et al., 2010; Heydrich and Blanke, 2013; Ronchi et al., 2015; Salomon et al., 2018), consisting of specialized substrates organized in a posterior to anterior structural progression, with posterior parts representing the primary cortical representations of interoceptive stimuli from contralateral body parts and more anterior parts, tested here in the right hemisphere, acting as integration areas for sensory signals and contextual cues ultimately leading to interoceptive awareness.

## Data availability

The data that support the findings of this study are available from the corresponding author upon reasonable request.

## Acknowledgements

We thank the all the stroke patients for their kindness and willingness to take part in the study, as well as the healthy participants. We are also particularly grateful to Sonia Ponzo, Amanda Hornsby, and Arturo Kerbel, for their help with patient recruitment and testing. No conflicts of interest were reported.

## Contributions

Sahba.B, L.C., A.F., P.M.J., L.P.K, and C.P. conceived and designed the experiment.

Sahba.B, L.C., L.P.K, V.M., and C.P. performed the experiment.

N.W. helped recruiting the stroke patients on his ward.

V.M. and Sara.B. drew the patients’ lesions.

L.P.K analyzed the data.

L.P.K wrote the manuscript, conjointly with A.F.

All authors read, corrected and approved the final manuscript.

## Footnotes

1 Normal t-tests were used as the behavioural data entered in the VLSM models were normally distributed (De Haan & Karnath, 2018).

## REFERENCES

Ackerley, R., Wasling, H. B., Liljencrantz, J., Olausson, H., Johnson, R. D., & Wessberg, J. (2014). Human C-tactile afferents are tuned to the temperature of a skin-stroking caress. Journal of Neuroscience, 34(8), 2879–2883.

Baier, B., Zu Eulenburg, P., Geber, C., Rohde, F., Rolke, R., Maihöfner, C., … & Dieterich, M. (2014). Insula and sensory insular cortex and somatosensory control in patients with insular stroke. European journal of pain, 18(10), 1385–1393.

Bates, E., Wilson, S. M., Saygin, A. P., Dick, F., Sereno, M. I., Knight, R. T., & Dronkers, N. F. (2003). Voxel-based lesion–symptom mapping. Nature neuroscience, 6(5), 448.

Björnsdotter, M. (2016). Brain Processing of CT-Targeted Stimulation. In Affective Touch and the Neurophysiology of CT Afferents (pp. 187–194). Snpringer, New York, NY.

Bisiach, E., Vallar, G., Perani, D., Papagno, C., & Berti, A. (1986). Unawareness of disease following lesions of the right hemisphere: anosognosia for hemiplegia and anosognosia for hemianopia. Neuropsychologia, 24(4), 471–482.

Case, L. K., Laubacher, C. M., Olausson, H., Wang, B., Spagnolo, P. A., & Bushnell, M. C. (2016). Encoding of touch intensity but not pleasantness in human primary somatosensory cortex. Journal of Neuroscience, 36(21), 5850–5860.

Case, L. K., Laubacher, C. M., Richards, E. A., Spagnolo, P. A., Olausson, H., & Bushnell, M. C. (2017). Inhibitory rTMS of secondary somatosensory cortex reduces intensity but not pleasantness of gentle touch. Neuroscience letters, 653, 84–91.

Catani, M., & de Schotten, M. T. (2012). Atlas of human brain connections. Oxford University Press.

Cauda, F., D’Agata, F., Sacco, K., Duca, S., Geminiani, G., & Vercelli, A. (2011). Functional connectivity of the insula in the resting brain. Neuroimage, 55(1), 8–23.

Craig, A. D., & Craig, A. D. (2009). How do you feel-now? The anterior insula and human awareness. Nature reviews neuroscience, 10(1).

Craig, A. D. (2010). The sentient self. Brain structure and function, 214, 563–577.

Critchley, H. D., Wiens, S., Rotshtein, P., Öhman, A., & Dolan, R. J. (2004). Neural systems supporting interoceptive awareness. Nature neuroscience, 7(2), 189.

Crucianelli, L., Metcalf, N. K., Fotopoulou, A. K., & Jenkinson, P. M. (2013). Bodily pleasure matters: velocity of touch modulates body ownership during the rubber hand illusion. Frontiers in psychology, 4, 703.

Crucianelli, L., Krahé, C., Jenkinson, P. M., & Fotopoulou, A. K. (2018). Interoceptive ingredients of body ownership: Affective touch and cardiac awareness in the rubber hand illusion. Cortex, 104, 180–192.

Curran-Everett, D. (2000). Multiple comparisons: philosophies and illustrations. American Journal of Physiology-Regulatory, Integrative and Comparative Physiology, 279(1), R1–R8.

de Haan, B., & Karnath, H. O. (2018). A hitchhiker’s guide to lesion-behaviour mapping. Neuropsychologia, 115, 5–16.

Dijkerman, H. C., & De Haan, E. H. (2007). Somatosensory processing subserving perception and action: Dissociations, interactions, and integration. Behavioral and brain sciences, 30(2), 224–230.

de Schotten, M. T., Bizzi, A., Dell’Acqua, F., Allin, M., Walshe, M., Murray, R., … & Catani, M. (2011). Atlasing location, asymmetry and inter-subject variability of white matter tracts in the human brain with MR diffusion tractography. Neuroimage, 54(1), 49–59.

Dubois, B., Slachevsky, A., Litvan, I., & Pillon, B. F. A. B. (2000). The FAB: a frontal assessment battery at bedside. Neurology, 55(11), 1621–1626.

Heydrich, L., & Blanke, O. (2013). Distinct illusory own-body perceptions caused by damage to posterior insula and extrastriate cortex. Brain, 136(3), 790–803.

Gentsch, A., Panagiotopoulou, E., & Fotopoulou, A. (2015). Active interpersonal touch gives rise to the social softness illusion. Current Biology, 25(18), 2392–2397.

Guarantors of Brain. London: W.B. Saunders (1986).

Khalsa, S. S., Rudrauf, D., Feinstein, J. S., & Tranel, D. (2009). The pathways of interoceptive awareness. Nature neuroscience, 12(12), 1494.

Kann, S., Zhang, S., Manza, P., Leung, H. C., & Li, C. S. R. (2016). Hemispheric lateralization of resting-state functional connectivity of the anterior insula: association with age, gender, and a novelty-seeking trait. Brain connectivity, 6(9), 724–734.

Kirsch, L. P., Krahé, C., Blom, N., Crucianelli, L., Moro, V., Jenkinson, P. M., & Fotopoulou, A. (2018). Reading the mind in the touch: Neurophysiological specificity in the communication of emotions by touch. Neuropsychologia, 116, 136–149.

Kurth, F., Zilles, K., Fox, P. T., Laird, A. R., & Eickhoff, S. B. (2010). A link between the systems: functional differentiation and integration within the human insula revealed by meta-analysis. Brain Structure and Function, 214(5-6), 519–534.

Lenoir, C., Algoet, M., & Mouraux, A. (2018). Deep continuous theta burst stimulation of the operculo-insular cortex selectively affects Aδ-fibre heat pain. The Journal of physiology, 596(19), 4767–4787.

Liew, S. L., Anglin, J. M., Banks, N. W., Sondag, M., Ito, K. L., Kim, H., … & Lefebvre, S. (2018). A large, open source dataset of stroke anatomical brain images and manual lesion segmentations. Scientific data, 5, 180011.

Lincoln, N. B., Jackson, J. M., & Adams, S. A. (1998). Reliability and revision of the Nottingham Sensory Assessment for stroke patients. Physiotherapy, 84(8), 358–365.

Löken, L. S., Wessberg, J., McGlone, F., & Olausson, H. (2009). Coding of pleasant touch by unmyelinated afferents in humans. Nature neuroscience, 12(5), 547.

Maier, O., Schröder, C., Forkert, N. D., Martinetz, T., & Handels, H. (2015). Classifiers for ischemic stroke lesion segmentation: a comparison study. PloS one, 10(12), e0145118.

McGlone, F., Wessberg, J., & Olausson, H. (2014). Discriminative and affective touch: sensing and feeling. Neuron, 82(4), 737–755.

McIntosh, R. D., Brodie, E. E., Beschin, N., & Robertson, I. H. (2000). Improving the clinical diagnosis of personal neglect: a reformulated comb and razor test. Cortex, 36(2), 289–292.

Mori, S., Wakana, S., Nagae-Poetscher, L. M., & Van Zijl, P. C. M. (2006). MRI atlas of human white matter. American Journal of Neuroradiology, 27(6), 1384.

Morrison, I. (2016). ALE meta-analysis reveals dissociable networks for affective and discriminative aspects of touch. Human brain mapping, 37(4), 1308–1320.

Nasreddine, Z. S., Phillips, N. A., Bédirian, V., Charbonneau, S., Whitehead, V., Collin, I., … & Chertkow, H. (2005). The Montreal Cognitive Assessment, MoCA: a brief screeningtool for mild cognitive impairment. Journal of the American Geriatrics Society, 53(4), 695–699.

Olausson, H., Lamarre, Y., Backlund, H., Morin, C., Wallin, B. G., Starck, G., … & Bushnell, M. C. (2002). Unmyelinated tactile afferents signal touch and project to insular cortex. Nature neuroscience, 5(9), 900.

Perini, I., & Olausson, H. (2015). Seeking pleasant touch: neural correlates of behavioral preferences for skin stroking. Frontiers in behavioral neuroscience, 9,

Preusser, S., Thiel, S. D., Rook, C., Roggenhofer, E., Kosatschek, A., Draganski, B., … & Pleger, B. (2014). The perception of touch and the ventral somatosensory pathway. Brain, 138(3), 540–548.

Robinson, R. G., Kubos, K. L., Starr, L. B., Rao, K., & Price, T. R. Mood disorders in stroke patients: importance of location of lesion. Brain, 107(1), 81–93.

Ronchi, R., Bello-Ruiz, J., Lukowska, M., Herbelin, B., Cabrilo, I., Schaller, K., & Blanke, O. (2015). Right insular damage decreases heartbeat awareness and alters cardio-visual effects on bodily self-consciousness. Neuropsychologia, 70, 11–20.

Rorden, C., & Brett, M. (2000). Stereotaxic display of brain lesions. Behavioural neurology, 12(4), 191–200.

Salomon, R., Ronchi, R., Dönz, J., Bello-Ruiz, J., Herbelin, B., Martet, R., … & Blanke, O. (2016). The insula mediates access to awareness of visual stimuli presented synchronously to the heartbeat. Journal of Neuroscience, 36(18), 5115–5127.

Salomon, R., Ronchi, R., Dönz, J., Bello-Ruiz, J., Herbelin, B., Faivre, N., … & Blanke, O. (2018). Insula mediates heartbeat related effects on visual consciousness. Cortex, 101, 87–95.

Triscoli, C., Olausson, H., Sailer, U., Ignell, H., & Croy, I. Frontiers in behavioral neuroscience 7, 208 (2013).

Tzourio-Mazoyer, N., Landeau, B., Papathanassiou, D., Crivello, F., Etard, O., Delcroix, N., … & Joliot, M. (2002). Automated anatomical labeling of activations in SPM using a macroscopic anatomical parcellation of the MNI MRI single-subject brain. Neuroimage, 15(1), 273–289.

Vocat, R., Staub, F., Stroppini, T., & Vuilleumier, P. (2010). Anosognosia for hemiplegia: a clinical-anatomical prospective study. Brain, 133(12), 3578–3597.

Wechsler, D. (1994). San Antonio, TX: Psychological Corporation; 1997. Wechsler abbreviated scale of intelligence.

Wechsler, D. (2001). Wechsler individual achievement test II-abbreviated. San Antonio, TX: The Psychological Corporation.

Whyte, E. M., & Mulsant, B. H. (2002). Post stroke depression: epidemiology, pathophysiology, and biological treatment. Biological psychiatry, 52(3), 253–264.

Wilson, B. A., Cockburn, J., & Halligan, P. W. Thames Valley Test Company.(1987b). Behavioural Inattention Test.

Zigmond, A., & Snaith, R. P. (1983). The hospital depression and anxiety scale. Acta Psychiatrica Scandinavica, 67(6), 361–70.

