## Supplementary material for "Damage to the Right Insula Disrupts the Perception of Affective Touch"

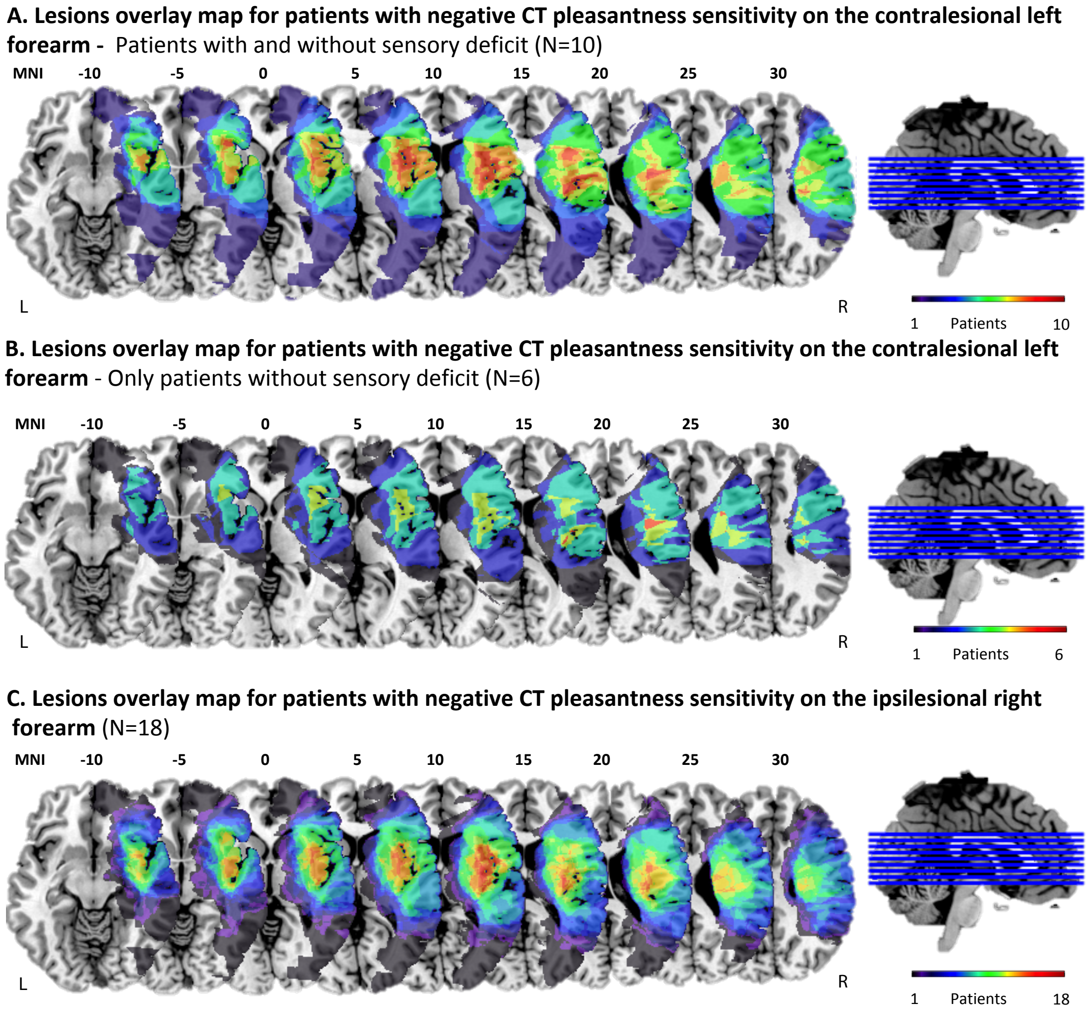


**Supplementary Figure 1.** A. Lesions overlap map for patients with negative CT pleasantness sensitivity on the left contralesional forearm, among all patients (N=10). B. Lesions overlap map for patients with negative CT pleasantness sensitivity on the left contralesional forearm, only in patients without sensory deficit (N=6). C. Lesions overlap map for patients with negative CT pleasantness sensitivity on the right ipsilesional forearm (N=18). An overlay heat map of participants' lesions was calculated from all lesions and superimposed on the chi2bet template brain using MRICron (Rorden et al., 2007).

**Supplementary Table 1. Number of significant voxels (from the atlas of grey matter – AAL – and white matter – JHU – and NatBrainLab's atlas) resulting from the VLSM analysis with the general pleasantness sensitivity scores** (velvet-sandpaper average pleasantness ratings), N=36**.**

| **Regions necessary for imagined tactile pleasantness sensitivity** | | | | | | |
|---|---|---|---|---|---|---|
| | **Region** | **Volume** | **x** | **y** | **z** | **T-value** |
| **AAL** | Unclassified | 111 | 24 | -18 | 6 | 3.23 |
| | Caudate | 48 | 13 | 14 | 4 | 2.85 |
| | Putamen | 14 | 21 | 4 | 8 | 2.69 |
| | Pallidum | 69 | 22 | -5 | 7 | 2.86 |
| | Thalamus | 19 | 20 | -13 | 7 | 2.96 |
| **JHU** | Unclassified | 22 | -5 | 7 | 22 | 2.86 |
| | Anterior_limb_of_int | 15 | 14 | 4 | 15 | 2.85 |
| | Posterior_limb_of_in | 24 | -18 | 6 | 24 | 3.23 |
| | Retrolenticular_part | 25 | -22 | 7 | 25 | 2.60 |
| | Posterior_corona_rad | 27 | -34 | 25 | 27 | 2.53 |
| **NatBrainLab** | Unclassified | 82 | 13 | 14 | 4 | 2.85 |
| | Corpus_Callosum | 1 | 22 | -28 | 28 | 2.53 |
| | Cortico_Ponto_Cerebellum | 6 | 19 | -10 | 12 | 2.49 |
| | Cortico_Spinal | 96 | 23 | -18 | 6 | 3.10 |
| | Internal_Capsule | 76 | 24 | -18 | 6 | 3.23 |
